# Opioid withdrawal produces sex-specific effects on fentanyl-vs.-food choice and mesolimbic transcription

**DOI:** 10.1101/2021.01.14.426662

**Authors:** E. Andrew Townsend, R. Kijoon Kim, Hannah L. Robinson, Samuel A. Marsh, Matthew L. Banks, Peter J. Hamilton

## Abstract

**Background:** Opioid withdrawal is a key driver of opioid addiction and an obstacle to recovery. However, withdrawal effects on opioid reinforcement and mesolimbic neuroadaptation are understudied and the role of sex is largely unknown.

**Methods:** Male (n=10) and female (n=9) rats responded under a fentanyl-vs.-food “choice” procedure during daily 2h sessions. In addition to the daily choice sessions, rats were provided extended access to fentanyl during 12h sessions. After two weeks of this self-administration regimen, the nucleus accumbens (NAc) and ventral tegmental area (VTA) of a subset of rats were subjected to RNA sequencing. In the remaining rats, a third week of this self-administration regimen was conducted, during which methadone effects on fentanyl-vs.-food choice were determined.

**Results:** Prior to opioid dependence, male and female rats similarly allocated responding between fentanyl and food. Abstinence from extended fentanyl access elicited a similar increase in somatic withdrawal signs in both sexes. Despite similar withdrawal signs and extended access fentanyl intake, opioid withdrawal was accompanied by a maladaptive increase in fentanyl choice in males, but not females. Behavioral sex differences corresponded with transcriptional hyperfunction in the NAc and VTA of opioid-withdrawn females relative to males. Methadone blocked withdrawal-associated increases in fentanyl choice in males, but failed to further decrease fentanyl choice in females.

**Conclusions:** These results provide foundational evidence of sex-specific neuroadaptations to opioid withdrawal, which may be relevant to the female-specific resilience to withdrawal-associated increases in opioid choice and aid in the identification of novel therapeutic targets.

## 1. Introduction

Opioid Use Disorder (OUD) diagnoses increased nearly 500% from 2010-2016 in the United States and continues to be a public health crisis (1). In response to this crisis, the United States Food and Drug Administration initiated a dialogue with OUD patients to identify critical barriers to recovery (2). Patients emphasized that opioid withdrawal was a primary driver of continued opioid use and relapse, with some participants describing themselves as “being a prisoner” to withdrawal during periods of opioid dependence (2). These patient reports highlight the role of opioid withdrawal in OUD, particularly as it relates to opioid-taking behaviors.

Consistent with these patient reports, opioid withdrawal enhances opioid self-administration in preclinical studies (3-8). Conversely, opioid withdrawal also decreases self-administration of non-opioid reinforcers such as food (3) and electrical brain stimulation (9). These findings suggest that opioid withdrawal can enhance opioid reinforcement not only through direct mechanisms, but also indirectly through the simultaneous devaluation of non-opioid reinforcers. Consistent with this hypothesis, opioid withdrawal increases the allocation of operant behavior (i.e., choice) emitted towards opioid injections over concurrently available palatable food in male nonhuman primates (10-14) and rats (15). Choice procedures such as these may be especially useful for studying the mechanisms of opioid withdrawal on both drug and non-drug reinforcement, as the maladaptive choice of drugs at the expense of non-drug alternatives is increasingly recognized as a hallmark of Substance Use Disorders (16-19).

The high efficacy mu-opioid receptor (MOR) agonist methadone is the most effective medication for OUD and is administered to patients with a high degree of opioid dependence (20). In contextual agreement with its clinical usage, methadone has little effect on opioid choice in non-dependent laboratory animals (13). However, in opioid-dependent non-human primates, methadone and other high efficacy MOR agonists block withdrawal-associated increases in opioid choice and promote behaviors maintained by alternative reinforcers (10-14, 21). Overall, the sensitivity of preclinical opioid-vs.-food choice procedures to MOR agonist treatments selectively in opioid-withdrawn subjects supports the translational utility of these procedures to study the mechanisms of withdrawal-induced behavioral misallocation.

Although sex differences in preclinical opioid abuse-related endpoints are well documented in non-opioid dependent subjects, less is known about opioid withdrawal affects sex differences in opioid reinforcement. Recent preclinical rodent studies suggest opioid withdrawal leads to protracted, sex-specific effects on somatic withdrawal signs, opioid self-administration, and reinstatement (e.g., (22-24)). However, sex differences in opioid-withdrawal effects on opioid-vs.-food choice are unexplored. We therefore examined sex differences in spontaneous opioid-withdrawal effects on fentanyl-vs.-food choice. We also performed RNA sequencing (RNAseq) in reward-associated brain regions of the nucleus accumbens (NAc) and ventral tegmental area (VTA) of opioid-withdrawn male and female rats as an unbiased and hypothesis-generating approach for evaluating the transcriptional consequences of opioid withdrawal in mesolimbic structures known to be involved in opioid reinforcement (25). Our results illustrate a sexually dimorphic behavioral response to fentanyl withdrawal, with withdrawal increasing fentanyl choice in males (a maladaptive behavioral allocation towards heightened drug use) and opioid withdrawal decreasing fentanyl choice in females (a pro-adaptive behavioral allocation towards food consumption). Sex differences in decision making corresponded with heightened transcription in the NAc and VTA of opioid-withdrawn females relative to males. In sum, this work demonstrates that female rats are resilient to the withdrawal-associated increases in fentanyl choice observed in males. Furthermore, the female-specific resilience to increased fentanyl choice may, in part, be the consequence of heightened mesolimbic transcriptional neuroadaptation.

## 2. Methods

### 2.1. Subjects

Twenty-seven Sprague-Dawley rats (14 male, 13 female) were acquired at 10 weeks of age (Envigo Laboratories, Frederick, MD) and implanted with vascular access ports (Instech, Plymouth Meeting, PA) and custom-made intravenous (i.v.) jugular catheters (26). Rats were administered subcutaneous (s.c.) ketoprofen (5 mg/kg) immediately and 24 h after surgery. Rats were singly housed and maintained on a 12-h light/dark cycle (lights off at 6:00 PM). Water and food (Teklad Rat Diet, Envigo) were provided ad-libitum in the home cage. Catheter patency was verified within 24-h of study conclusion by instantaneous muscle tone loss following intravenous (i.v.) methohexital (1.6 mg) administration. Animal maintenance and research were conducted in accordance with the 2011 guidelines for the care and use of laboratory animals and protocols approved by the Virginia Commonwealth University Institutional Animal Care and Use Committee.

### 2.2. Apparatus and Catheter Maintenance

Operant chambers located in sound-attenuating cubicles (Med Associates, St. Albans, VT) were equipped with two retractable levers, a set of three LED lights (red, yellow, green) mounted above each lever, and a retractable “dipper” cup (0.1 ml) located between the levers for presenting diluted Ensure® (18% or 32% v/v vanilla flavor Ensure® in tap water; Abbott Laboratories, Chicago, IL). Drug infusions were delivered by a syringe pump (PHM-100, Med Associates) located inside the cubicle (27). After each behavioral session, catheters were flushed with gentamicin (0.4 mg) and 0.1 ml of heparinized saline (10 U/ml).

### 2.3. Drugs

Fentanyl HCl was provided by the National Institute on Drug Abuse Drug Supply Program (Bethesda, MD). (±)-Methadone HCl was purchased from Spectrum Chemicals (Gardena, CA). Methohexital sodium was purchased from the Virginia Commonwealth University pharmacy (Richmond, VA). Drugs were dissolved in sterile water and diluted with sterile saline. Solutions were passed through a 0.22-micron sterile filter (Millex GV, Millipore Sigma, Burlington, MA) before administration. Drug doses were expressed as the salt forms and delivered based on bodyweight.

### 2.4. Procedure

#### 2.4.1. Self-administration Training

##### 2.4.1.1. Fentanyl-vs.-food choice training

Rats (10 male, 9 female) were trained to respond under a fentanyl-vs.-food choice procedure as described previously (28, 29) and detailed in the *Supplemental Materials*. Briefly, each choice session consisted of five, 20-min response components wherein the available unit fentanyl dose increased between each component and the food reinforcer remained constant (18% Ensure®). Food availability was signaled by illuminating a red stimulus light above the left lever and fentanyl availability was signaled by illuminating a green stimulus light above the right lever (longer flashes signaled availability of larger unit fentanyl doses). During each component, rats could complete up to 10 ratio requirements (i.e., fixed-ratio (FR) 5) between the food- and drug-associated levers. Choice was considered stable when the smallest fentanyl unit dose that maintained at least 80% of completed ratio requirements on the drug-associated lever was within a 0.5 log unit for three consecutive days with no trends (i.e., stability criteria). Stability criteria were not assessed until after at least five choice sessions. Data collected during the final training day served as “baseline” values for subsequent analyses.

##### 2.4.1.2. Saline-vs.-food choice training

Eight rats (4 male, 4 female) were trained to respond under a saline-vs.-food choice procedure. Rats were trained to self-administer 32% v/v Ensure® under a FR5, TO20 schedule using training criteria described in *Supplemental Materials*. The Ensure® concentration was increased from 18% to 32% to accelerate acquisition. Rats were then provided access to i.v. saline (5s infusion / 315g). Saline was available under a FR1, TO20 for two sessions and FR5, TO20 for three sessions. Finally, saline and 32% Ensure® were made available under the same choice parameters and stability criteria as described above for fentanyl.

#### 2.4.2. Extended access self-administration

Following stability, a regimen of overnight (6:00 PM − 6:00 AM) extended access (FR5, TO10 schedule of reinforcement) to fentanyl (3.2 μg/kg/injection) or saline (5s injection / 315g) began. 8h after each extended access self-administration session (∼1:55 PM), rats were observed for 30s for the presence of nine somatic withdrawal signs (see *Supplemental Materials*) and weighed before beginning the daily choice test (2:00 PM). Overnight self-administration tests occurred Sunday-Thursday nights for two consecutive weeks (i.e., Week 1 and Week 2) in all rats, although a subset of rats progressed to a third week (see below).

### 2.5. Tissue Preparation and RNA Sequencing

A representative sample of the fentanyl-trained rats (4 male, 4 female; **Supplemental Figure 1**) and all of the saline-trained rats (4 male, 4 female; **Supplemental Figure 2**) were used in RNA sequencing (RNAseq) studies. In lieu of the final choice session (2:00 P.M. ±1h), rats were rapidly decapitated without anesthesia, brains were sectioned into 1 mm coronal slices using sex-specific brain matrices (Zivic Instruments: Pittsburgh, PA; Male: BSRLS001-1; Female: BSRLA001-1), and bilateral tissue punches were collected from the nucleus accumbens (NAc; 12 gauge; internal diameter 2.16 mm) and ventral tegmental area (VTA; 16 gauge; internal diameter 1.19 mm). Tissue was frozen on dry ice and stored at −80°C. RNA was extracted and purified (RNeasy, Qiagen) according to manufacturer’s instructions. Total RNA was quantified with the Qubit RNA HS Assay Kit (Thermo Fisher). RNA quality control assays were performed on the TapeStation 4200 (Agilent), and the RNA Integrity Number (RIN) for all samples ranged from 8.0 to 9.2 (mean value 8.7±0.05). Ribosomal RNA depletion and library preparation (Illumina Ribo-Zero) was performed and RNAseq was carried out at Genewiz with the following configuration: 2×150 paired-end reads on an Illumina sequencing platform (HiSeq 2500) with a sequencing depth of ∼22M reads per sample (mean value 22±0.03M). Other overall sample sequencing statistics include the Mean Quality Score (35.54±0.03) and the percent of bases ≥ 30 (91.94±0.16).

Sequence reads were trimmed to remove possible adapter sequences and nucleotides with poor quality using Trimmomatic v.0.36. The trimmed reads were mapped to the Rattus norvegicus Rnor6.0 reference genome available on ENSEMBL using the STAR aligner v.2.5.2b. Unique gene hit counts were calculated by using featureCounts from the Subread package v.1.5.2. After extraction of gene hit counts, the gene hit counts table was used for downstream differential expression analysis. Using DESeq2, a comparison of gene expression between groups of samples was performed. The Wald test was used to generate p-values and log2-fold changes. Genes with an adjusted *p*-value < 0.05 and absolute log2-fold change > 1 were called as differentially expressed genes for each comparison. For pattern identification in union heat maps, venn diagrams, and gene ontology, genes with an unadjusted *p*-value < 0.05 and absolute log2 fold change > 1.3 were called as differentially expressed genes for each comparison. Heatmaps were generated using Morpheus and gene ontology analysis was performed using GOrilla, running all ontologies (Process, Function, Component) and reporting uncorrected enrichment *p-*values (30).

### 2.6. Methadone Tests

Six male and five female fentanyl-trained rats progressed to a third week of extended fentanyl access. Somatic withdrawal signs, weights, and fentanyl-vs.-food choice testing continued as in Weeks 1 and 2. Rats were administered counterbalanced s.c. pretreatments of saline or methadone (0.1, 0.32, 1, 3.2 mg/kg) 10 min prior to fentanyl-vs.-food choice tests.

### 2.7. Data Analysis

Behavioral data were analyzed using Prism 8 (GraphPad, La Jolla, CA) and Geisser-Greenhouse corrections for sphericity when appropriate. Choice-related dependent measures were analyzed by two-way ANOVA (or a mixed-effects analysis in instances of missing data) or by paired t-tests. Other dependent measures include injections earned per 12 hours and bodyweight. These dependent measures were analyzed within each sex by one-way ANOVA and between sex by two-way ANOVA. Withdrawal signs were analyzed by non-parametric Friedman tests.

Significant (*p*<0.05) interactions were followed by Dunnett, Sidak, or Dunn post-hoc tests as appropriate. See **Supplemental Table S1** for a complete reporting of statistical results.

## 3. Results

### 3.1. Fentanyl-vs.-food choice in non-dependent rats

At “baseline”, liquid food was almost exclusively chosen when no fentanyl or small unit doses (0.32, 1 μg/kg/injection) were available (dashed lines; **Figure 1D**; **Figure 2A, 2B**). As the fentanyl dose increased, behavior was reallocated towards the fentanyl lever and the largest unit doses (3.2, 10 μg/kg/injection) maintained near exclusive choice. Additionally, choices per component decreased as a function of increasing fentanyl doses (dashed lines; **Figure 1E**; **Figure 2C, 2D**). No baseline sex differences were detected for either choice-related measure.

**Figure 1.**
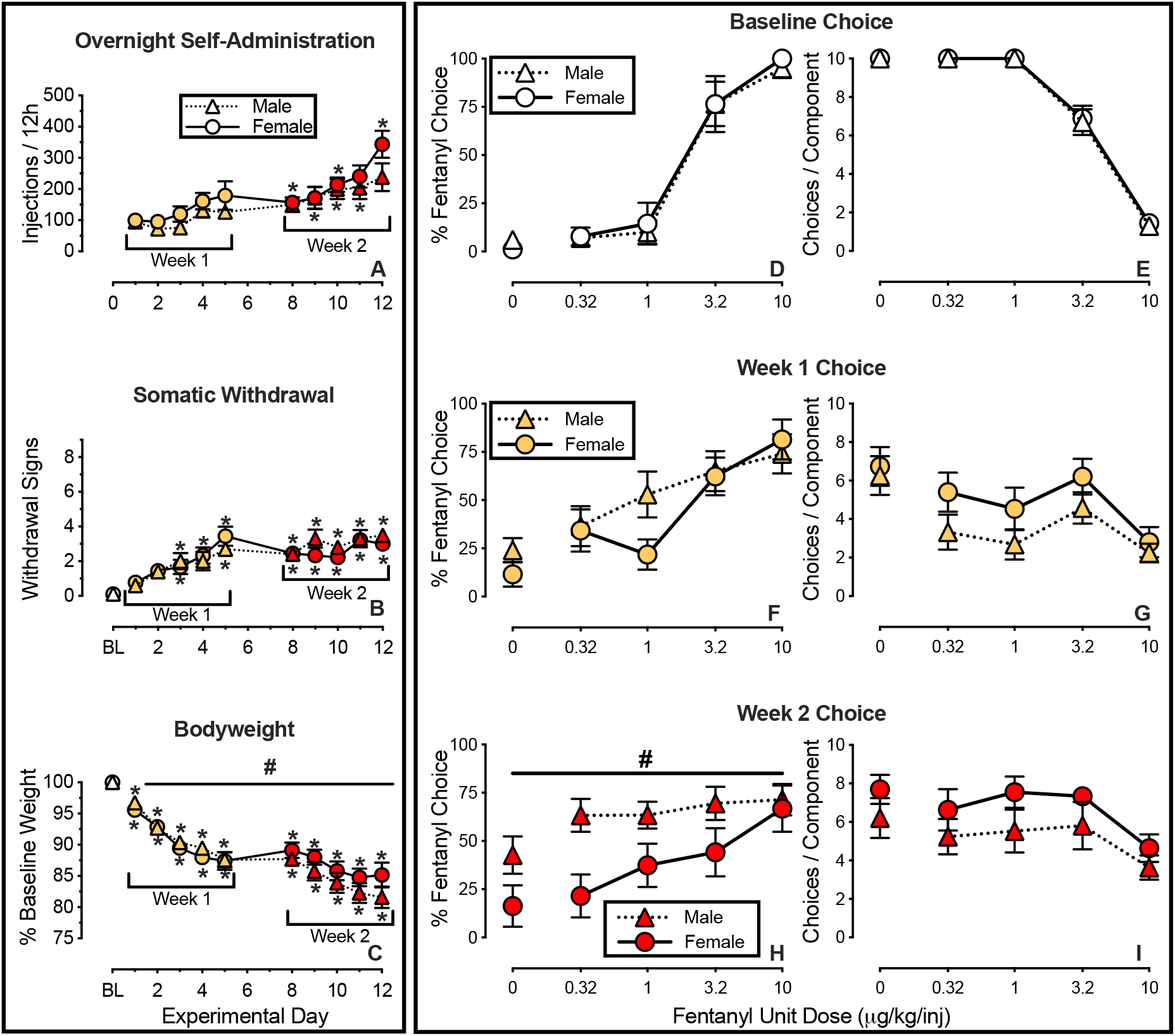
Opioid withdrawal promotes sex-specific effects on fentanyl-vs.-food choice: Decreased fentanyl choice in female relative to male rats. Left panels (A-C): Escalating self-administration of fentanyl and its effects on somatic withdrawal sign expression and bodyweight in male (n=10, dotted line) and female (n=9, solid line) rats. Abscissae of left panels (A-C): experimental day. Top ordinate (A): number of fentanyl injections (3.2 μg/kg unit dose) during each 12-h session. Middle ordinate (B): number of somatic withdrawal signs (maximum of nine) observed 8h after the conclusion of each overnight self-administration session. Bottom ordinate (C) change in bodyweight expressed as a percentage of baseline. Right panels (D-I): Sex comparison of opioid-withdrawal effects on fentanyl-vs.-food choice assessed 8h after overnight (6PM-6AM) fentanyl self-administration. Abscissae: intravenous unit fentanyl dose in μg/kg/injection. Left ordinates (D, F, H): percentage of completed ratio requirements on the fentanyl-associated lever. Right ordinates (E, G, I): number of choices completed per component. Points represent mean ± SEM. * Denotes significant difference relative to Day 1 (A) or Baseline (B-C), which are placed above the symbol for female rats and below the symbol for female rats. **#** Denotes a significant main effect of sex. Significance defined as *p* < 0.05. See *Supplemental Table 1* for statistics relevant to each panel.

**Figure 2.**
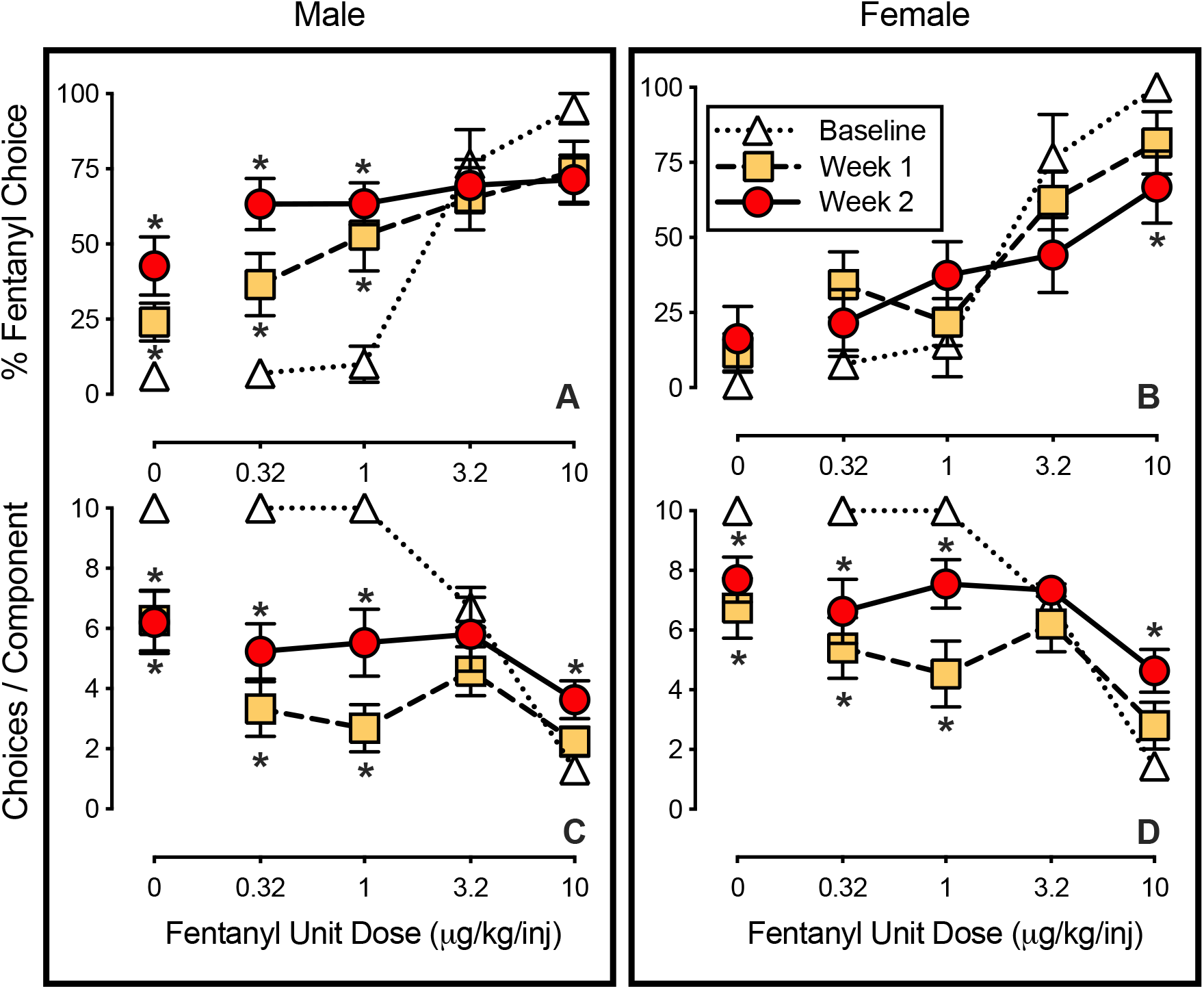
Withdrawal from overnight fentanyl self-administration produces opposing, sex-specific effects on fentanyl-vs.-food choice. Fentanyl-vs.-food choice results assessed 8h after overnight (6PM-6AM) fentanyl self-administration sessions in male (n=10, left) and female (n=9, right) rats. Abscissae: intravenous unit fentanyl dose in μg/kg/injection. Top ordinates (A, B): percentage of completed ratio requirements on the fentanyl-associated lever. Bottom ordinates (C, D): number of choices completed per component. Points represent mean ± SEM. * Denotes a significant difference at a unit fentanyl dose relative to Baseline. Significance defined as *p* < 0.05. See *Supplemental Table 1* for statistics relevant to each panel.

### 3.2. Withdrawal from overnight fentanyl self-administration produces opposing, sex-specific effects on fentanyl-vs.-food choice

Fentanyl self-administration escalated across two weeks of extended access (**Figure 1A**, male: F_1.5,13.8_=12.2, *p*=0.002; female: F_2.2,17.3_=7.7, *p*=0.003). Over this same time, somatic withdrawal signs increased (**Figure 1B**, male: Friedman statistic=54.2, *p*<0.0001; female: Friedman statistic=46.7, *p*<0.0001), and body weights decreased (**Figure 1B**, male: F_2.2,19.5_=83.3, *p*<0.0001; female: F_2.5,20.2_=31.0, *p*<0.0001). Other than greater bodyweight decreases in males (sex × time interaction: F_10,170_=2.3, *p*=0.02), no other sex differences were detected for these dependent measures.

Opioid withdrawal resulted in sex-dependent effects on fentanyl-vs.-food choice. Whereas percent fentanyl choice was similar between sexes at baseline (**Figure 1D**) and during the first week of extended fentanyl access (**Figure 1F**), fentanyl choice was significantly lower in females relative to males during the second week of extended fentanyl access (**Figure 1H**: main effect of sex F_1,17_=5.6, *p*=0.03). The number of choices completed per session did not differ between sexes.

Analyses of withdrawal effects *within* each sex revealed opposing effects on fentanyl choice. Whereas percent fentanyl choice progressively increased across Weeks 1 and 2 in males (**Figure 2A**: interaction: F_3.6,30_=9.5, *p*<0.0001), fentanyl choice in females was unaffected by opioid withdrawal at Week 1 and slightly, but significantly, *decreased* by Week 2 (**Figure 1B**: interaction: F_4.3,33.6_=4.3, *p*=0.006). Opioid withdrawal similarly decreased the number of choices completed per component within each sex (**Figure 1C** (male): interaction: F_3.7,33.3_=15.2, *p*<0.0001; **Figure 1D** (female): interaction: F_3.8,30.4_=9.5, *p*<0.0001).

### 3.3. Mesolimbic transcriptional consequences of opioid withdrawal

Sex-matched rats subjected to a saline-vs.-food choice paradigm were used as reference tissue to generate differentially expressed genes (DEGs) in fentanyl-withdrawn rats. In transcriptional analyses from the NAc, females demonstrated a far greater diversity and degree of up- and down-regulated transcripts than males (**Figure 3A-B**). By reducing statistical thresholds to enable broad pattern recognition and organizing the female transcripts from down to up-regulated in a union heatmap, nearly no concordance in the direction and degree of transcript expression was observed across males and females (**Figure 3C**). Further, while the NAc of females possessed a far greater number of up- and down-regulated transcripts than males, only approximately 6% of the female-identified transcripts were similarly regulated in males (**Figure 3D-E**). Lastly, we explored the broad gene ontology (GO) terms enriched in sex-specific up- and down-regulated transcript lists. In male-specific up-regulated transcripts, the top GO terms were enriched for the biological processes of protein kinase activity and signal transduction (**Figure 3F**). The top GO terms in female-specific up-regulated transcripts were enriched in the cytosol and linked to the functions of oxygen transport and pre-miRNA binding (**Figure 3G**). No significantly enriched GO terms were identified in the male down-regulated transcripts. The female-specific down-regulated transcripts revealed top GO terms linked to the neuronal component, specifically the glutamatergic synapse, as well as cellular energetics and protein phosphorylation (**Figure 3H**).

**Figure 3.**
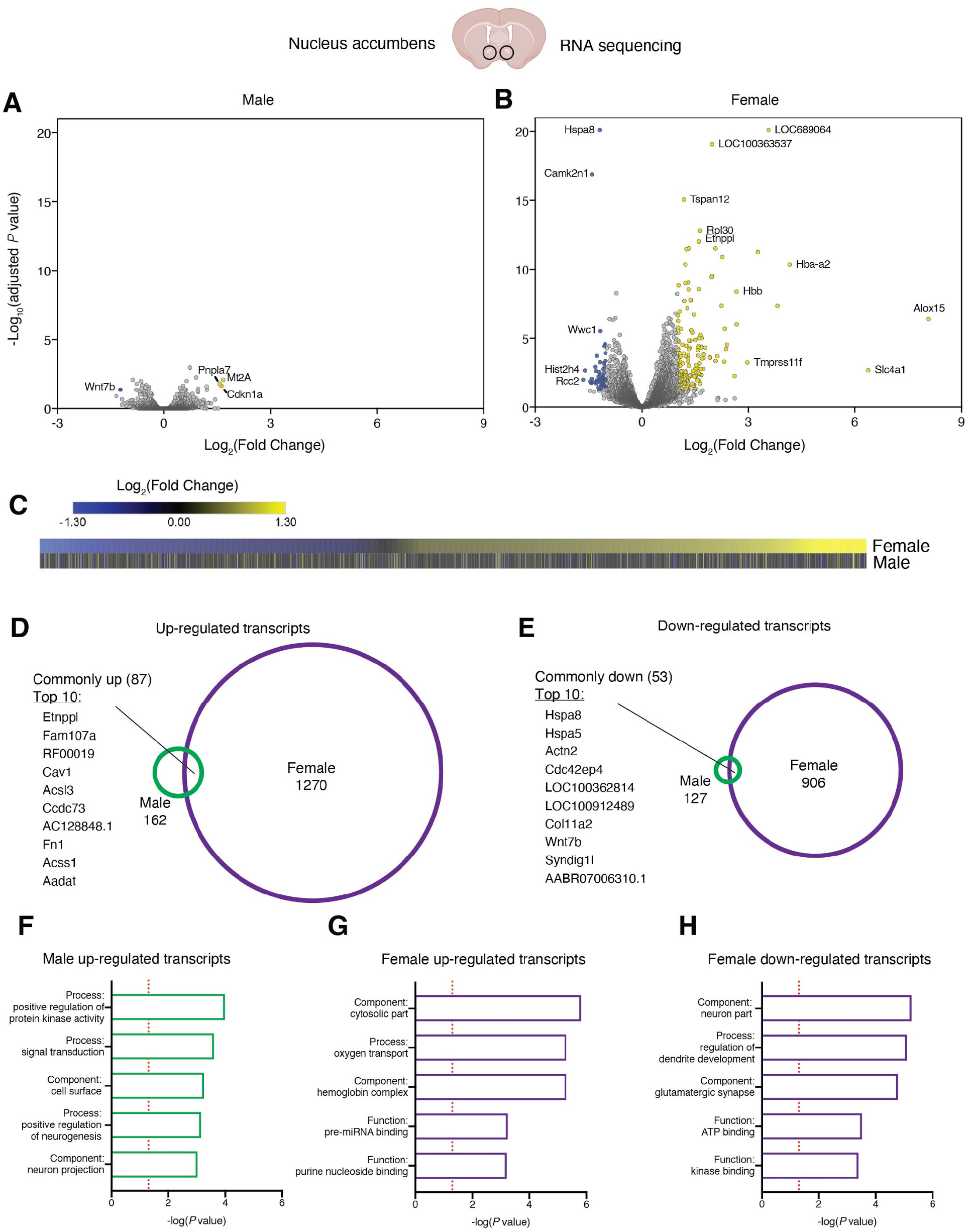
Female and male rats experience distinct transcriptional activity within the nucleus accumbens following repeated fentanyl withdrawal. Volcano plots depicting the differentially expressed genes (DEGs) from microdissected nucleus accumbens identified in male (n = 4) (A) and female (n = 4) (B) rats. DEGs were normalized to sex-matched rats subjected to saline-vs.-food protocol. DEG cutoff for volcano plots are Wald test adjusted *p* value < 0.05 and absolute log2 fold change > 1. (C) Union heatmap comparing female transcripts organized from down-regulated to up-regulated (top) to male transcripts of the same identity (bottom). Significance cutoffs for this and subsequent analyses raw *p* value < 0.05 and absolute log2 fold change > 1.3, to enable broad pattern recognition. Venn diagrams depicting the degree of overlap in up-regulated (D) and down-regulated (E) transcripts across males and females. (F-H) GO terms enriched within the sex-specific regions of the Venn diagrams.

Similar results were observed within the VTA. Again, we observed that females experienced more transcripts regulated to a greater degree than males (**Figure 4A-B**), minimal concordance in the identity and degree of regulation specific transcripts across sexes (**Figure 4C**), and only modest overlap in which transcripts are up- or down-regulated across males and females (**Figure 4D-E**). In sex-specific GO analysis, we identified that neuronal projection and enzymatic activity related terms were enriched in the male up-regulated transcripts (**Figure 4F**). Like in the NAc, female up-regulated transcripts were linked to oxygen carrier activity (**Figure 4G**). Relatively fewer GO terms reached significance for male and female down-regulated transcripts, with males associated with lipid processing (**Figure 4H**) and females associated with regulation of cellular secretion (**Figure 4I**).

**Figure 4.**
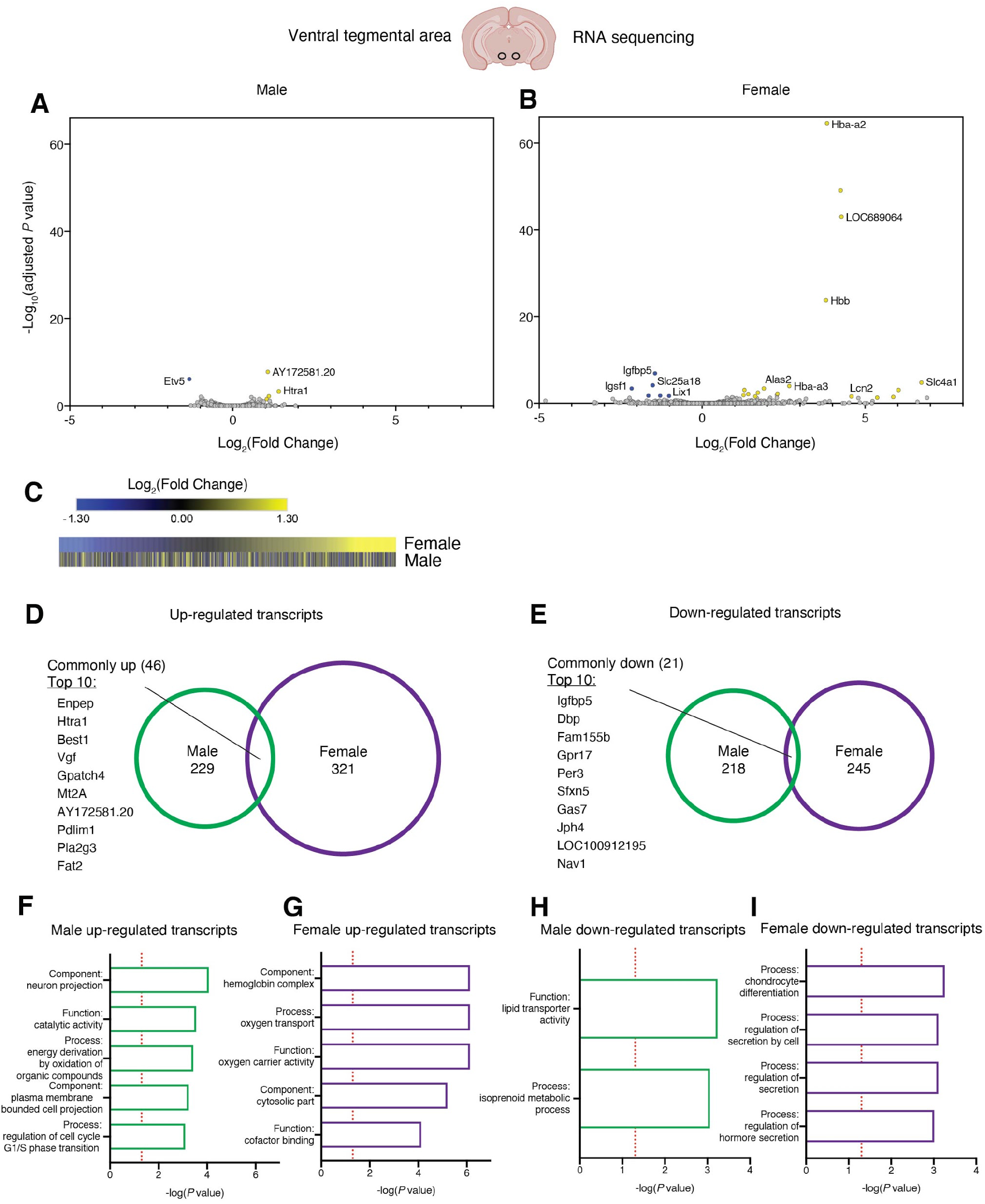
Female and male rats experience distinct transcriptional activity within the ventral tegmental area following repeated fentanyl withdrawal. Volcano plots depicting the differentially expressed genes (DEGs) from microdissected ventral tegmental area identified in male (n = 3) (A) and female (n = 3) (B) rats. DEGs were normalized to sex-matched rats subjected to saline-vs.-food protocol. DEG cutoff for volcano plots are Wald test adjusted *p* value < 0.05 and absolute log2 fold change > 1. (C) Union heatmap comparing female down- and up-regulated transcripts (top) to male transcripts (bottom). Significance cutoffs for this and subsequent analyses raw *p* value < 0.05 and absolute log2 fold change > 1.3, to enable broad pattern recognition. Degree of overlap in up-regulated (D) and down-regulated (E) transcripts across males and females. (F-I) GO terms enriched within the sex-specific regions of the venn diagrams.

In comparing the transcriptional responses within sex and across the NAc and VTA, males had a similar number and minimal overlap in their up- and down-regulated transcripts across the NAc and VTA (**Supplemental Figure 3A**). In contrast, females had far more affected transcripts in the NAc and a greater degree of overlapping affected transcripts across these brain regions (**Supplemental Figure 3B**).

### 3.5. Acute methadone decreases fentanyl-vs.-food choice in opioid-withdrawn male, but not female, rats

A subset of rats were subjected to a third week of extended fentanyl self-administration and choice testing (**Figure 5A**), and saline or methadone injections preceded each choice session. In male rats, 3.2 mg/kg methadone decreased percent session fentanyl choice relative to saline (**Figure 5B**: T=2.9, df=5, *p*=0.03) and increased the overall number of food choices (**Figure 5C**: interaction: F_1.9,9.3_=53.5, *p*<0.0001). Methadone-induced decreases in fentanyl choice did not correspond with decreased somatic withdrawal signs (**Supplemental Figure 4E**). In female rats, 3.2 mg/kg methadone did not significantly alter percent session fentanyl choice (**Figure 5B**), the number of choices completed per session (**Figure 5C**), or somatic withdrawal signs (**Supplemental Figure 4F**). The effectiveness of smaller methadone doses are shown in **Supplemental Figure 4**.

**Figure 5.**
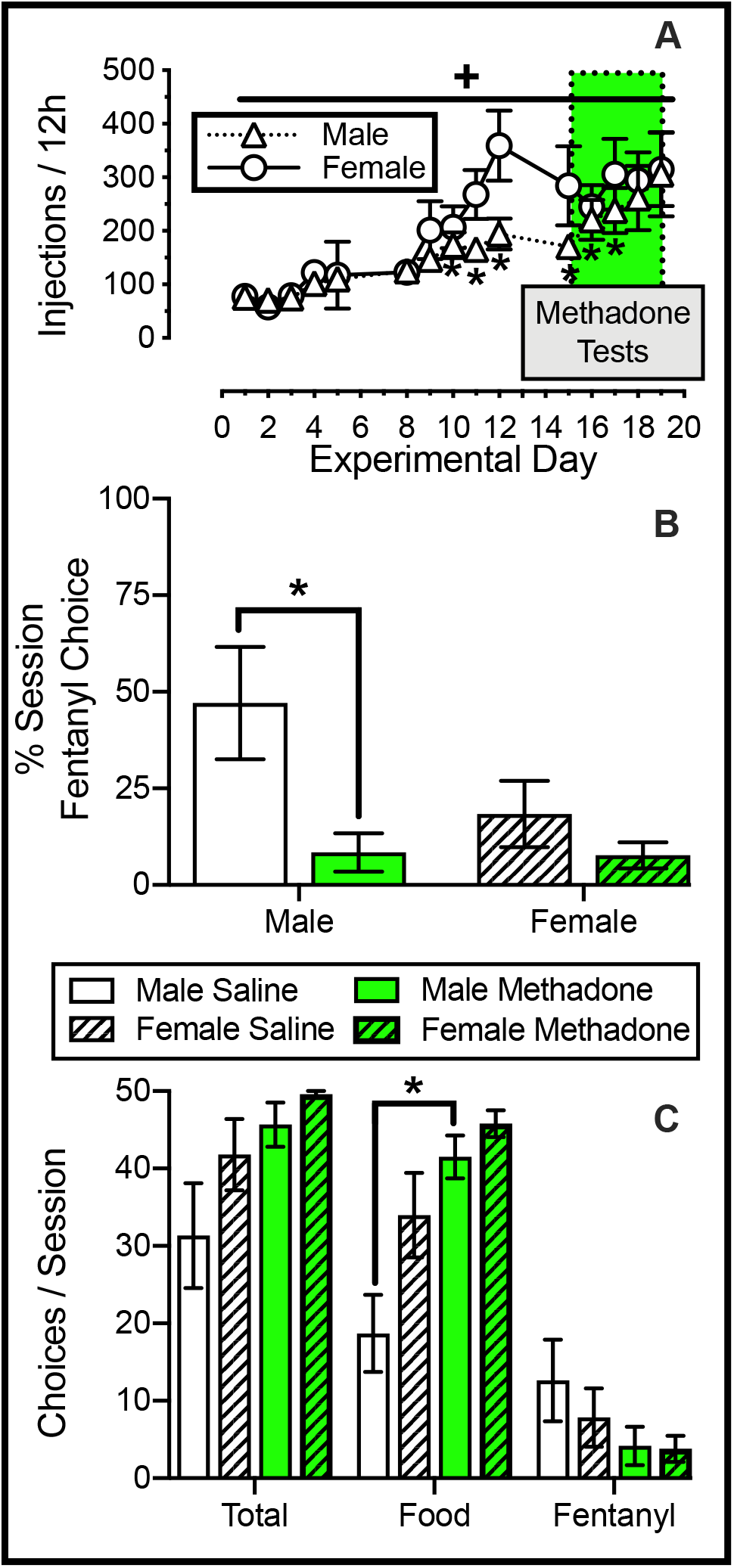
Acute methadone decreases fentanyl choice in male, but not female, opioid-withdrawn rats. Effects of acute methadone treatment on fentanyl choice in male (n=6) and female (n=5) rats during Week 3 of overnight (6PM-6AM) fentanyl self-administration. Top abscissae: experimental day. Middle abscissae: treatment condition. Bottom abscissae: reinforcer type. Top ordinate (A): number of fentanyl injections (3.2 μg/kg unit dose) during each 12-h session. Middle ordinate (B): percentage of completed ratio requirements on the fentanyl-associated lever across the entire session. Bottom ordinate: (C): number of choices completed per session. Points represent mean ± SEM. * Denotes significant difference at a unit fentanyl dose relative to Day 1 in panel A. **+** Denotes a main effect of time in panel A. *Denotes difference from saline within a sex in panels B-C. Significance defined as *p* < 0.05. See *Supplemental Table 1* for statistics relevant to each panel.

## 4. Discussion

Here we identify sexually dimorphic decision-making strategies in an opioid-vs.-food choice context following opioid withdrawal: Whereas opioid withdrawal increased choice of fentanyl over a food alternative in males, fentanyl choice slightly decreased in opioid-withdrawn female rats. The female-specific capacity to spontaneously allocate effort towards a pro-adaptive food reinforcer while undergoing withdrawal mirrors the effects of methadone on opioid choice in opioid-withdrawn males within this procedure, and suggests the neural substrates of the female-specific behavior allocation within the context of drug-choice may aid in the identification of novel therapeutic targets for OUD. To this end, we performed RNAseq profiling of mesolimbic brain areas, revealing striking transcriptional hyperactivity in the NAc and VTA of females relative to males and involved predominantly non-overlapping transcript identities across sexes. These results provide foundational evidence of sex-specific neuroadaptations to repeated opioid withdrawal, which may be relevant to the female-specific resilience to withdrawal-associated increases in opioid choice.

Fentanyl intake similarly escalated in male and female rats (**Figure 1A**). The present results are consistent with previous opioid escalation studies in male rats (31). However, the present results are in contrast to reports of female rodents showing greater rates of opioid escalation than males (24, 32-34) in some studies, but not all (35-37). Reasons for inconsistent sex differences in rates of opioid self-administration are not presently clear and could reflect differences in either the self-administered opioid or other experimental conditions. Nevertheless, the spontaneous expression of somatic withdrawal signs was similar between sexes (**Figure 1B**), suggesting that sex differences in fentanyl choice and mesolimbic transcription were not attributable to differences in opioid intake.

Non-opioid dependent male and female rats similarly allocated choice between fentanyl injections and a concurrently available food reinforcer (**Figure 1D**). However, spontaneous opioid withdrawal unmasked behavioral sex differences (**Figure 1H**). Opioid choice robustly increased in male opioid-withdrawn rats (**Figure 2A**), consistent with decades of preclinical studies using male opioid-withdrawn subjects (10-15). Conversely, opioid withdrawal coincided with a small, but significant, decrease in opioid choice in females (**Figure 2B**), representing the first evaluation of opioid-vs.-food choice in female opioid-withdrawn subjects. Consistent with the present results, a recent study reported rates of oxycodone self-administration to be decreased in female, but not male, rats during protracted opioid withdrawal (23). Collectively, these findings illustrate an adaptive response to opioid withdrawal in female rats, which promotes choice of a non-drug alternative over continued opioid self-administration.

By performing unbiased transcriptional profiling of mesolimbic brain regions, we aimed to identify the brain-region specific molecular correlates of this sex-divergent phenotype. Surprisingly, we observed that the NAc and VTA of females was more transcriptionally active than the corresponding male brain regions. The female NAc was the region of the most substantial transcriptional activity (**Supplemental Figure 3**), hinting that the aggregate consequence of transcription in this brain region may largely contribute to the observed female-specific behavior to allocate effort towards non-opioid reinforcers while in withdrawal, possibly through altered reward processing. Further, we observed minimal overlap in transcript identities between males and females in either brain region. This suggests that the male and female mesolimbic neuroadaptations in response to opioid withdrawal were almost entirely distinct. Also notable was the greater agreement of similarly regulated individual transcripts between the NAc and VTA of females when compared to males, suggesting a greater degree of co-regulation across mesolimbic brain regions in females.

Male and female transcriptional adaptations in the NAc and VTA may predominantly occur within neuronal cell-types, as revealed by our GO analysis enriched for component terms associated with the neuron and neuron projection component (**Figure 3F, H, Figure 4F**). In the NAc, male up-regulated transcripts were linked to an increase in protein kinase activity (**Figure 3F**), whereas female down-regulated transcripts were enriched for kinase binding GO terms (**Figure 3H**). This raises the interesting possibility that NAc protein kinase function was broadly increased in males yet decreased in females, which would have divergent consequences on NAc function, and may partially explain the observed sex-specific behaviors. Moreover, within females, the top GO terms were nearly identical for up-regulated transcripts across the NAc (**Figure 3G**) and VTA (**Figure 4G**). These were enriched with GO terms relating to hemoglobin-mediated oxygen transport, driven by some of the most strongly up-regulated genes in both brain areas: Hba-a2, LOC689064, Hbb, and Hba-a3. Similar GO terms were not observed for female-specific down-regulated transcripts across the NAc (**Figure 3H**) and VTA (**Figure 4I**). Finally, an interesting observation was that many of these down-regulated female transcripts within the NAc were related to neuronal dendrite development, glutamatergic synaptic neurotransmission, and cellular energetics (**Figure 3H**) whereas down-regulated female transcripts within the VTA were related to the GO functions of secretion (**Figure 4I**). This hints that the net consequence of female-specific transcriptional down-regulation converges on diminished glutamatergic pre- and post-synaptic neurotransmission within the NAc as well as diminished neurotransmitter release emanating from the VTA. It is possible, therefore, that the observed female-specific transcriptional dynamics were an adaptive, active response to opioid withdrawal which consequently resulted in an altered function of NAc cell-types, granting females the capacity to allocate effort towards pro-adaptive reinforcers while in opioid withdrawal. How these transcriptional responses relate to cellular function will be an important area for future research.

Methadone decreased fentanyl choice in opioid-withdrawn male, but not female, rats (**Figure 5B**). The results in male rats are in agreement with previous studies in male non-human primates showing that MOR agonists reverse opioid withdrawal-associated increases in opioid choice (10-14, 21). Furthermore, these results are in line with the clinical benefits of methadone, which include not only decreasing illicit opioid use (20, 38, 39), but also increasing quality of life through engagement with non-opioid reinforcers such as social relationships and leisure activities (40, 41). Interestingly, the methadone-induced decrease in fentanyl choice in male rats was similar to the spontaneous decrease in fentanyl choice observed in opioid-withdrawn female rats. An intriguing future direction is whether chronic administration of efficacious OUD pharmacotherapies (e.g., methadone) to opioid dependent male rats induces similar mesolimbic transcriptional hyperactivity as observed in untreated, opioid-withdrawn female rats.

The failure of methadone to decrease fentanyl choice in female rats is seemingly inconsistent with the clinical literature, wherein methadone is more effective to promote long-term abstinence from illicit opioids in women than in men (42, 43). The lack of methadone effects in opioid-withdrawn female rats is most likely attributable to lower fentanyl choice in the absence of methadone in females relative to males (**Figure 5B**: males: 47%; females: 18%), providing a smaller window to detect methadone-induced decreases in fentanyl choice in female rats. Indeed, 3.2 mg/kg methadone decreased session fentanyl choice to approximately 8% in males and females alike. Overall, these data contribute to emerging preclinical data showing sex-specific effects of methadone in opioid-withdrawn rats (44).

The current study extended the study of opioid-withdrawal effects on opioid choice to include sex as a biological variable. Opioid withdrawal robustly increased opioid choice in male subjects, which aligns with the prominent role of opioid withdrawal in maintaining maladaptive opioid use in OUD patients (2). Our finding of decreased opioid choice in female rats does not appear to have a clinical correlate, as women are clearly negatively affected by OUD (45).

However, the current findings in female rats may reflect species-specific responses to opioid withdrawal. Alternatively, these results may suggest that opioid withdrawal has a lesser influence on continued opioid use in women than in men, with other factors contributing to OUD severity. Nonetheless, female rats may serve as a model organism for a phenotype resilient to withdrawal-associated increases in opioid choice. Here we used RNAseq to begin exploring neurobiological distinctions between male and female opioid-withdrawn rats, with the ultimate goal of identifying targetable substrates to diminish the ability of opioid withdrawal to bias decision-making processes towards increased opioid use at the expense of non-opioid alternatives.

