## Supplemental Materials for "Opioid withdrawal produces sex-specific effects on fentanyl-vs.-food choice and mesolimbic transcription"

***Training of fentanyl and Ensure® self-administration***

Rats were first trained to respond on the right lever for IV fentanyl (n=10 male, 9 female; 3.2 µg/kg/infusion) under a fixed-ratio 1, 20s timeout (FR1, TO20) schedule of reinforcement, signaled by the illumination of a green stimulus light. After rats earned at least ten fentanyl infusions under the FR1, TO20 schedule, the response requirement was increased to FR5. This FR5, TO20 schedule was in place for at least five sessions and until the number of earned infusions was stable, defined as the number of earned injections differing by less than 20% of the running mean for three consecutive sessions with no upward or downward trends. Next, rats were trained to respond on the left lever for a 5 s presentation of 18% Ensure® under a FR1, TO20 schedule of reinforcement, signaled by the illumination of a red stimulus light. After rats earned at least fifty food presentations under the FR1, TO20 schedule, the response requirement for food presentations was increased to FR5. The FR5, TO20 schedule of food reinforcement was in place until three consecutive days with at least 50 earned reinforcers and with no upward- or downward-facing trends.

Next, fentanyl was made concurrently available with 18% Ensure® under a FR5, TO20: FR5, TO20 schedule of reinforcement. Here, the behavioral session consisted of five 20-min response components wherein the concentration of the food reinforcer was held constant and the available unit dose of fentanyl increased with each of the five response components. Each response component was preceded by a "sample" component, consisting of a non-contingent infusion of the unit fentanyl dose available during the upcoming response component followed by a 2-min time out. Next, a 5-s presentation of the liquid-food dipper was programmed followed by a 2-min time out. Following this second time out, the response component began. During each response component, both levers extended, a red stimulus light above the left lever was illuminated to signal liquid food availability and a green stimulus light above the right lever illuminated to signal i.v. fentanyl availability. Response requirement (FR5) completion on the left lever resulted in a 5-s presentation of the food dipper whereas response requirement (FR5) completion on the right lever resulted in the delivery of the i.v. fentanyl dose available for that component. Responding on one lever reset the ratio requirement for the other lever. A different drug dose was available during each of the five successive response components. Specifically, the unit fentanyl doses were as follows: (0 (no injection), 0.32, 1.0, 3.2, and 10 µg/kg/infusion during components 1–5, respectively). Fentanyl doses were varied by changing the infusion duration (e.g., 315 g rat: 0, 0.5, 1.56, 5, and 15.6 s during components 1–5, respectively) and the green light above the drug-associated lever flashed on and off in 3-s cycles (i.e., longer flashes corresponded with larger fentanyl doses).

During each response component, rats could complete up to 10 total ratio requirements between the food- and drug-associated levers. Each ratio requirement completion initiated a 20 s time out, the retraction of both levers, and darkening of the red and green stimulus lights. If all 10 ratio requirements were completed before 20 min had elapsed, then both levers retracted, and stimulus lights were extinguished for the remainder of that component. Choice was considered stable when the smallest fentanyl unit dose that maintained at least 80% of completed ratio requirements on the drug-associated lever was within a 0.5 log unit for three consecutive days with no trends (i.e., stability criteria). During the training phase, stability criteria were not assessed until rats responded in at least 5 choice sessions. Data collected during the final day of training were used as "baseline" values for subsequent analyses.

***Scoring of Somatic Withdrawal Signs***

Somatic withdrawal signs were assessed approximately 8h after the conclusion of each overnight fentanyl or saline self-administration session (~1:55 PM). Here, the observer (EAT) visually inspected each rat inside of its clear plastic home cage for 30s. If a rat was found to be in a rest/sleep posture at the outset of the observation period, the observer would awaken the rat with three taps on the outside of the cage. During this observation period, the presence of absence of nine somatic withdrawal signs (1) were recorded: 1) Paw Tremor: a transient raising of one of the front paws accompanied by shaking of that paw; 2) Teeth Chatter: a very rapid opening a closing of the jaw; 3) Eye Twitch: a spastic movement of both eyes inwards and outwards from the socket; 4) Mastication: a chewing motion that is not directed towards eating, drinking, or grooming; 5) Yawn: a compound behavior of opening the jaw and closing of the eyes; 6) Wet Dog Shake: a whole-body movement that resembles an exaggerated shivering response; 7) Piloerection: bristling of fur that is typically matted close to the skin (e.g., facial region, dorsal area); 8) Diarrhea: amorphous feces present either in the bedding or on the rat; 9) Ptosis: a drooping of the upper eyelid for at least 1/3 of the observation period (10s).

**Supplemental Table 1. Statistical details (GraphPad Prism Version 8)**

| **Figure number** | **Factor name** | **F-value** | ***p*-value** |
|---|---|---|---|
| Figure 1A: Escalation One-way ANOVAs & Two-way ANOVA (sex) | **Male Time**, Dunnett's Post Hoc vs Night 1 | $\mathbf{F_{1.5,13.8}=12.2}$ | **0.002** |
| | **Female Time**, Dunnett's Post Hoc vs Night 1 | $\mathbf{F_{2.2,17.3}=7.7}$ | **0.003** |
| | **Time** | $\mathbf{F_{2.2,36}=17.7}$ | **<0.0001** |
| | Sex | $F_{1,17}=1.3$ | 0.28 |
| | Interaction | $F_{9,153}=1.0$ | 0.4 |
| Figure 1B: Withdrawal Signs Friedman Tests; Two-way ANOVA (sex) | **Male Time, Dunn's Post Hoc vs Baseline** | **Friedman stat=54.2** | **<0.0001** |
| | **Female Time, Dunn's Post Hoc vs Baseline** | **Friedman stat=46.7** | **<0.0001** |
| | **Time** | $\mathbf{F_{5.2,92.9}=11.3}$ | **<0.001** |
| | Sex | $F_{1,18}=0.13$ | 0.7 |
| | Interaction | $F_{9,162}=1.0$ | 0.5 |

| | | | |
|---|---|---|---|
| Figure 1C: Bodyweight One-way ANOVAs & Two-way ANOVA (sex) | **Male Time** (raw weights), Dunnett's Post Hoc vs Baseline<br>**Female Time** (raw weights), Dunnett's Post Hoc vs Baseline<br><br>**Time**<br>Sex (normalized weights)<br>**Interaction, Sidak Post Hoc** | **$F_{2.2,19.5}$=83.3**<br>**$F_{2.5,20.2}$=31.0**<br><br>**$F_{2.3,38.4}$=88.1**<br>$F_{1,17}$=0.3<br>**$F_{10,170}$=2.3** | **<0.0001**<br>**<0.0001**<br><br>**<0.001**<br>0.57<br>**0.02** |
| Figure 1D: Baseline % fentanyl choice (between sexes) Two-way ANOVA | **Fentanyl Unit Dose**<br>Sex<br>Interaction | **$F_{2.0,34.5}$=77.4**<br>$F_{1,17}$=0.04<br>$F_{4,68}$=0.16 | **<0.0001**<br>0.85<br>0.96 |
| Figure 1F: Week 1 % fentanyl choice (between sexes) Two-way ANOVA | **Fentanyl Unit Dose**<br>Sex<br>Interaction | **$F_{3.1,48.9}$=23.6**<br>$F_{1,17}$=0.5<br>$F_{4,63}$=2.5 | **<0.0001**<br>0.49<br>0.053 |
| Figure 1H: Week 2 % fentanyl choice (between sexes) Two-way ANOVA | **Fentanyl Unit Dose**<br>**Sex**<br>Interaction | **$F_{2.7,43.8}$=9.0**<br>**$F_{1,17}$=5.6**<br>$F_{4,66}$=1.8 | **0.0002**<br>**0.03**<br>0.15 |
| Figure 1E: Baseline # choices (between sexes) Two-way ANOVA | **Fentanyl Unit Dose**<br>Sex<br>Interaction | **$F_{1.1,18.6}$=321**<br>$F_{1,17}$=0.10<br>$F_{4,68}$=0.05 | **<0.0001**<br>0.76<br>0.99 |
| Figure 1G: Week 1 # choices (between sexes) Two-way ANOVA | **Fentanyl Unit Dose**<br>Sex<br>Interaction | **$F_{3.2,54.7}$=13.3**<br>$F_{1,17}$=1.7<br>$F_{4,68}$=0.79 | **<0.0001**<br>0.21<br>0.54 |
| Figure 1I: Week 2 # choices (between sexes) Two-way ANOVA | **Fentanyl Unit Dose**<br>Sex<br>Interaction | **$F_{2.8,48.4}$=7.0**<br>$F_{1,17}$=2.0<br>$F_{4,68}$=0.19 | **0.0006**<br>0.18<br>0.95 |
| Figure 2A: Withdrawal effects on % fentanyl choice (Male) Mixed Effects Analysis | **Fentanyl Unit Dose**<br>**Time**<br>**Interaction,** Dunnett's Post Hoc vs Baseline | **$F_{2.3,20.6}$=26.8**<br>**$F_{1.9,17.1}$=8.1**<br>**$F_{3.6,30}$=9.5** | **<0.0001**<br>**0.004**<br>**<0.0001** |
| Figure 2B: Withdrawal effects on % fentanyl choice (Female) Mixed Effects Analysis | **Fentanyl Unit Dose**<br>Time<br>**Interaction,** Dunnett's Post Hoc vs Baseline | **$F_{2.2,17.4}$=47.5**<br>$F_{1.5,12}$=0.23<br>**$F_{4.3,33.6}$=4.3** | **<0.0001**<br>0.73<br>**0.006** |
| Figure 2C: Withdrawal effects on # choices (Male) Two-way ANOVA | **Fentanyl Unit Dose**<br>**Time**<br>**Interaction,** Dunnett's Post Hoc vs Baseline | **$F_{2.9,26}$=32**<br>**$F_{1.7,15.3}$=13.7**<br>**$F_{3.7,33.3}$=15.2** | **<0.0001**<br>**0.0006**<br>**<0.0001** |
| Figure 2D: Withdrawal effects on # choices (Female) Two-way ANOVA | **Fentanyl Unit Dose**<br>**Time**<br>**Interaction,** Dunnett's Post Hoc vs Baseline | **$F_{2.7,21.7}$=57.7**<br>**$F_{1.5,11.9}$=9.9**<br>**$F_{3.8,30.4}$=9.5** | **<0.0001**<br>**0.005**<br>**<0.0001** |
| Figure 2E: Withdrawal effects on summary choice (Male) Two-way ANOVA | **Dependent measure**<br>**Time**<br>**Interaction,** Dunnett's Post Hoc vs Baseline | **$F_{1.3,11.3}$=32.7**<br>**$F_{1.9,16.8}$=10.2**<br>**$F_{2.6,23.5}$=29.6** | **<0.0001**<br>**0.0015**<br>**<0.0001** |
| Figure 2F: Withdrawal effects on summary choice (Female) Two-way ANOVA | **Dependent measure**<br>**Time**<br>**Interaction,** Dunnett's Post Hoc vs Baseline | **$F_{1.1,8.5}$=28.0**<br>**$F_{1.5,12}$=9.8**<br>**$F_{1.8,14.4}$=3.9** | **0.0005**<br>**0.005**<br>**0.05** |

| | | | |
|---|---|---|---|
| Figure 5A: Escalation One-way ANOVAs & Two-way ANOVA (sex) | **Male: Time, Dunnett's Post Hoc vs Night 1**<br><br>**Female: Time,** Dunnett's Post Hoc vs Night 1<br><br>**Time**<br>Sex<br>Interaction | **$F_{1.6,8.1}=9.2$**<br><br>**$F_{2.5,9.8}=9.8$**<br><br>**$F_{2.6, 23.7}=16$**<br>$F_{1,9}=1.1$<br>$F_{14,126}=1.5$ | **0.01**<br><br>**0.008**<br><br>**<0.0001**<br>0.3<br>0.13 |
| Figure 5B: Methadone Summary % Choice Paired t-tests & Two-way ANOVA (sex) | **Male Methadone**<br><br>Female Methadone<br><br>**Methadone**<br>Sex<br>Interaction | **T=2.9, df=5**<br><br>T=1.6, df=4<br><br>**$F_{1,9}=9.5$**<br>$F_{1,9}=1.9$<br>$F_{1,9}=3.0$ | **0.03**<br><br>**0.2**<br><br>**0.01**<br>0.2<br>0.1 |
| Figure 5C: Methadone Summary # Choices Separate two-way ANOVAs & Two-way ANOVA (sex) | **Male: Dependent measure**<br>**Time**<br>**Interaction**, Sidak Post Hoc<br><br>**Dependent measure**<br>Time<br>Interaction<br><br>**Dependent measure**<br>Sex<br>Interaction | **$F_{1.4,6.9}=22.1$**<br>**$F_{1,5}=9.6$**<br>**$F_{1.9,9.3}=53.5$**<br><br>**$F_{1.1,4.5}=68.7$**<br>$F_{1,4}=2.9$<br>$F_{1.1,4.5}=4.4$<br><br>**$F_{2.1,18.5}=52.0$**<br>$F_{1,9}=1.7$<br>$F_{5,45}=2.1$ | **0.002**<br>**0.03**<br>**<0.0001**<br><br>**0.0006**<br>0.16<br>0.10<br><br>**<0.0001**<br>0.23<br>0.08 |
| Supplemental Figure 1A: Fentanyl RNAseq Escalation One-way ANOVAs & Two-way ANOVA (sex) | Male Time (Fentanyl)<br>Female Time (Fentanyl)<br><br>**Time**<br>Sex<br>Interaction | $F_{1.3,4.0}=4$<br>$F_{1.5,4.6}=2.1$<br><br>**$F_{1.9,11.5}=4.8$**<br>$F_{1,6}=.13$<br>$F_{9,54}=1.1$ | 0.11<br>0.22<br><br>**0.03**<br>0.73<br>0.4 |
| Supplemental Figure 1B: Fentanyl RNAseq Withdrawal Signs Friedman Test; Two-way ANOVA (sex) | **Male (non-parametric Friedman Test),** Dunn's Post Hoc<br>**Female (non-parametric Friedman Test),** Dunn's Post Hoc<br><br>**Time**<br>Sex<br>Interaction | **Friedman stat=23.6**<br>**Friedman stat=24.0**<br><br>**$F_{2.7,16.3}=4.0$**<br>$F_{1,6}=0.003$<br>$F_{9,54}=0.81$ | **0.009**<br>**0.008**<br><br>**0.03**<br>0.96<br>0.61 |
| Supplemental Figure 1C: Fentanyl RNAseq Bodyweight One-way ANOVAs & Two-way ANOVA (sex) | **Male Time** (raw weights), Dunnett's Post Hoc vs Baseline<br>**Female Time** (raw weights), Dunnett's Post Hoc vs Baseline<br><br>**Time**<br>Sex (normalized weights)<br>**Interaction, Sidak Post Hoc** | **$F_{1.7,5.1}=74.2$**<br>**$F_{1.5,4.4}=23.2$**<br><br>**$F_{1.5,9.2}=80.2$**<br>$F_{1,6}=1.7$<br>**$F_{9,54}=3.9$** | **0.0002**<br>**0.006**<br><br>**<0.001**<br>0.24<br>**0.0008** |
| Supplemental Figure 1D: Fentanyl RNAseq Baseline % fentanyl choice (between sexes) Two-way ANOVA | **Fentanyl Unit Dose**<br>Sex<br>Interaction | **$F_{2.1,12.3}=112.2$**<br>$F_{1,6}=0.19$<br>$F_{4,24}=0.84$ | **<0.0001**<br>0.68<br>0.51 |
| Supplemental Figure 1F: Fentanyl RNAseq Week 1 % fentanyl choice (between sexes) Two-way ANOVA | **Fentanyl Unit Dose**<br>Sex<br>Interaction | **$F_{1.7,8.9}=11.2$**<br>$F_{1,6}=2.9$<br>$F_{4,21}=2.4$ | **0.005**<br>0.14<br>0.08 |
| Supplemental Figure 1H: Fentanyl RNAseq Week 2 % fentanyl choice (between sexes) Two-way ANOVA | **Fentanyl Unit Dose**<br>Sex<br>Interaction | **$F_{2.3,13.4}=7.8$**<br>$F_{1,6}=0.8$<br>$F_{4,23}=1.3$ | **0.004**<br>0.4<br>0.3 |

| | | | |
|---|---|---|---|
| Supplemental Figure 1E: Fentanyl RNAseq Baseline # choices (between sexes) Two-way ANOVA | **Fentanyl Unit Dose**<br>Sex<br>Interaction | **$F_{1.2,7.4}$=547.7**<br>$F_{1,6}$=0.11<br>$F_{4,24}$=0.11 | **<0.0001**<br>0.75<br>0.97 |
| Supplemental Figure 1G: Fentanyl RNAseq Week 1 # choices (between sexes) Two-way ANOVA | **Fentanyl Unit Dose**<br>Sex<br>Interaction | **$F_{2.3,13.6}$=4.9**<br>$F_{1,6}$=0.42<br>$F_{4,24}$=0.1 | **0.02**<br>0.54<br>0.98 |
| Supplemental Figure 1I: Fentanyl RNAseq Week 2 # choices (between sexes) Two-way ANOVA | **Fentanyl Unit Dose**<br>Sex<br>Interaction | **$F_{2.4,14.1}$=5.8**<br>$F_{1,6}$=0.09<br>$F_{4,24}$=0.82 | **0.01**<br>0.77<br>0.52 |
| Supplemental Figure 2A: Saline RNAseq Escalation One-way ANOVAs & Two-way ANOVA (sex) | Male Time (saline)<br>Female Time (Saline)<br><br>**Time**<br>Sex<br>Interaction | $F_{1.0,3.1}$=5.9<br>$F_{1.1,3.3}$=3.2<br><br>**$F_{1.1,6.6}$=8.3**<br>$F_{1,6}$=0.04<br>$F_{9,54}$=0.13 | 0.09<br>0.16<br><br>**0.02**<br>0.84<br>0.99 |
| Supplemental Figure 2B: Saline RNAseq Withdrawal Signs Friedman Test; Two-way ANOVA (sex) | Male (non-parametric Friedman Test)<br>Female (non-parametric Friedman Test)<br><br>Time<br>Sex<br>Interaction | Friedman stat=11.3<br>Friedman stat=15.3<br><br>$F_{2.2,13.1}$=2.5<br>$F_{1,6}$=0.86<br>$F_{9,54}$=0.26 | 0.33<br>0.12<br><br>0.12<br>0.39<br>0.98 |
| Supplemental Figure 2C: Saline RNAseq Bodyweight One-way ANOVAs & Two-way ANOVA (sex) | **Male Time** (raw weights), Dunnett's Post Hoc vs Baseline<br>Female Time (raw weights)<br><br>**Time**<br>**Sex (normalized weights)**<br>**Interaction, Sidak Post Hoc** | **$F_{2.4,7.3}$=23.9**<br>$F_{1.5,4.5}$=2.1<br><br>**$F_{2.2,13.5}$=9.5**<br>**$F_{1,6}$=14.8**<br>**$F_{9,54}$=5.3** | **0.0005**<br>0.22<br><br>**0.002**<br>**0.008**<br>**<0.0001** |
| Supplemental Figure 2D: Saline RNAseq Baseline % saline choice (between sexes) Two-way ANOVA | Component<br>Sex<br>Interaction | $F_{+infinity,+infinity}$=NA<br>$F_{1,28}$=NA<br>$F_{4,28}$=NA | NA<br>NA<br>NA |
| Supplemental Figure 2F: Saline RNAseq Week 1 % saline choice (between sexes) Two-way ANOVA | **Component**<br>Sex<br>Interaction | $F_{+infinity,+infinity}$=NA<br>$F_{1,28}$=NA<br>$F_{4,28}$=NA | NA<br>NA<br>NA |
| Supplemental Figure 2H: Saline RNAseq Week 2 % saline choice (between sexes) Two-way ANOVA | **Component**<br>Sex<br>Interaction | $F_{+infinity,+infinity}$=NA<br>$F_{1,28}$=NA<br>$F_{4,28}$=NA | NA<br>NA<br>NA |
| Supplemental Figure 2E: Saline RNAseq Baseline # choices (between sexes) Two-way ANOVA | Component<br>Sex<br>Interaction | $F_{4,24}$=0.46<br>$F_{1,6}$=1.7<br>$F_{4,24}$=0.46 | 0.39<br>0.24<br>0.77 |
| Supplemental Figure 2G: Saline RNAseq Week 1 # choices | Component<br>Sex<br>Interaction | $F_{1.4,8.6}$=1.9<br>$F_{1,6}$=0.05<br>$F_{4,24}$=1.5 | 0.21<br>0.82<br>0.22 |

| | | | |
|---|---|---|---|
| (between sexes) Two-way ANOVA | | | |
| Supplemental Figure 2I: Saline RNAseq Week 2 # choices (between sexes) Two-way ANOVA | Component<br>Sex<br>Interaction | $F_{1.9,11.6}=1.5$<br>$F_{1,6}=2.9$<br>$F_{4,24}=1.5$ | 0.26<br>0.14<br>0.23 |
| Supplemental Figure 4A: Male Methadone effect on Week 3 % Choice (Session) One-way ANOVA | **Treatment,** Dunnett's Post Hoc vs Saline | $\mathbf{F_{1.9,9.6}=4.7}$ | **0.04** |
| Supplemental Figure 4B: Female Methadone effect on Week 3 % Choice (Session) One-way ANOVA | Treatment | $F_{1.6,6.4}=2.8$ | 0.14 |
| Supplemental Figure 4C: Methadone effects on Summary Choice (Male) Two-way ANOVA | **Dependent measure**<br>Time<br>**Interaction,** Dunnett Post Hoc | $\mathbf{F_{1.9,9.4}=17.23}$<br>$F_{1.8,9.1}=3.7$<br>$\mathbf{F_{2.0,9.9}=7.3}$ | **0.0008**<br>0.07<br>**0.01** |
| Supplemental Figure 4D: Methadone effects on Summary Choice (Female) Two-way ANOVA | **Dependent measure**<br>Time<br>Interaction | $\mathbf{F_{1.2,5.0}=38.8}$<br>$F_{1.3,5.1}=0.3$<br>$F_{1.1,4.5}=3.4$ | **0.0013**<br>0.28<br>0.13 |
| Supplemental Figure 4E: Methadone effects on withdrawal signs (Male) Friedman Test | **Treatment (non-parametric Friedman Test),** Dunn's Post Hoc | **Friedman Statistic=11.4** | **0.004** |
| Supplemental Figure 4F: Methadone effects on withdrawal signs (Female) Friedman Test | Treatment (non-parametric Friedman Test) | Friedman Statistic=1.9 | 0.65 |

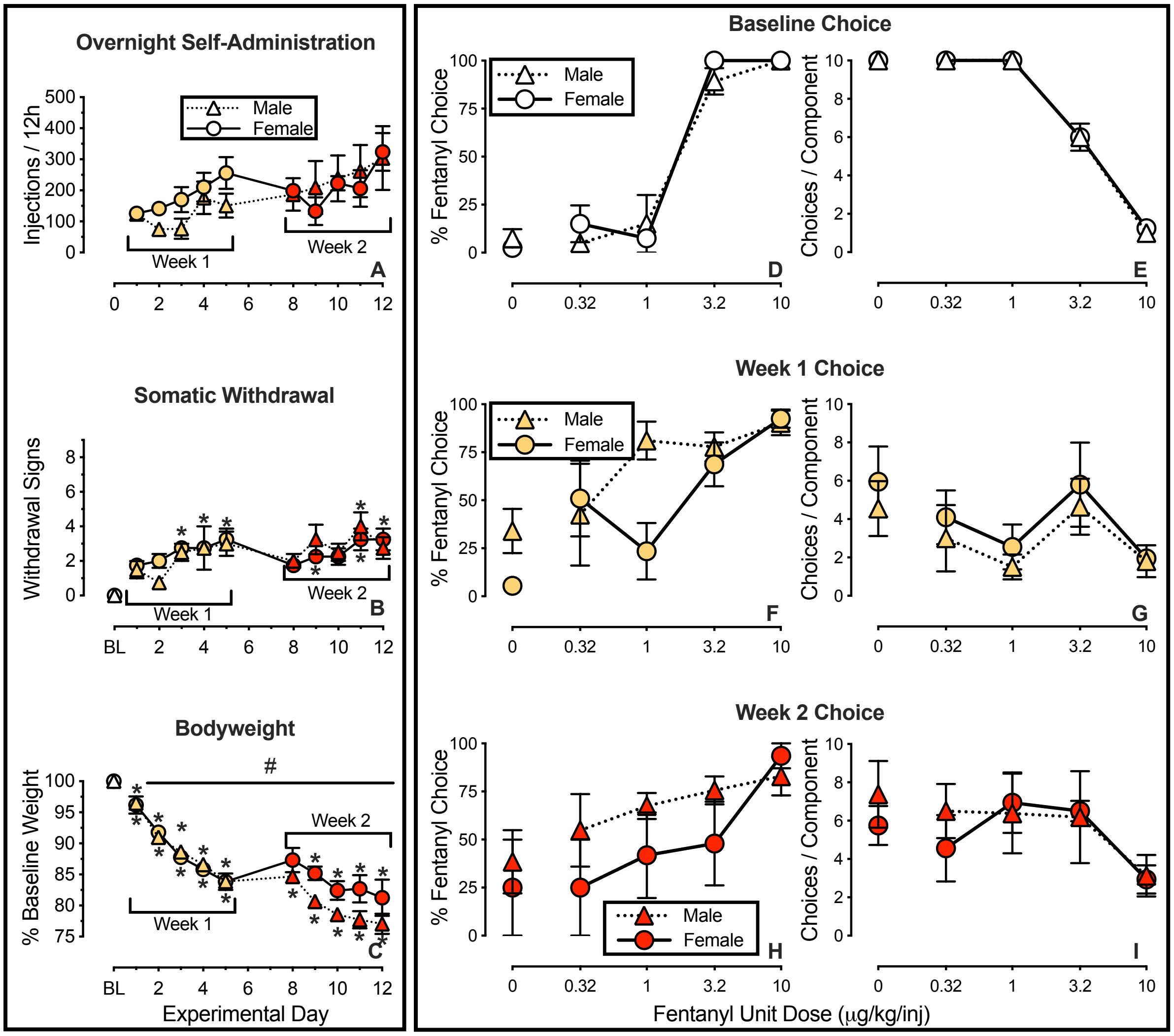

Supplemental Figure 1.

**Fentanyl self-administration and withdrawal-related metrics in rats used in RNA sequencing studies.** Left panels (A-C): Overnight (6AM-6PM) self-administration of fentanyl and its effects on somatic withdrawal sign expression and bodyweight in male (n=4, left) and female (n=4, right) rats. Abscissae of left panels (A-C): experimental day. Top ordinate (A): number of fentanyl infusions (3.2 µg/kg unit dose) during each 12-h session. Middle ordinate (B): number of somatic withdrawal signs (maximum of nine) observed 8h after the conclusion of each overnight self-administration session. Bottom ordinate (C) change in bodyweight expressed as a percentage of baseline. Right panels (D-I): Sex comparison of opioid-withdrawal effects on fentanyl-versus-food choice assessed 8h after overnight (6PM-6AM) fentanyl self-administration. Abscissae: intravenous unit fentanyl dose in µg/kg/infusion. Left ordinates (D, F, H): percentage of completed ratio requirements on the fentanyl-associated lever. Right ordinates (E, G, I): number of choices completed per component. Points represent mean ± SEM. * Denotes significant difference relative to Day 1 (A) or Baseline (B-C), which are placed above the symbol for female rats and below the symbol for female rats. # Denotes a significant main effect of sex. Significance defined as $p < 0.05$. See *Supplemental Table 1* for statistics relevant to each panel.

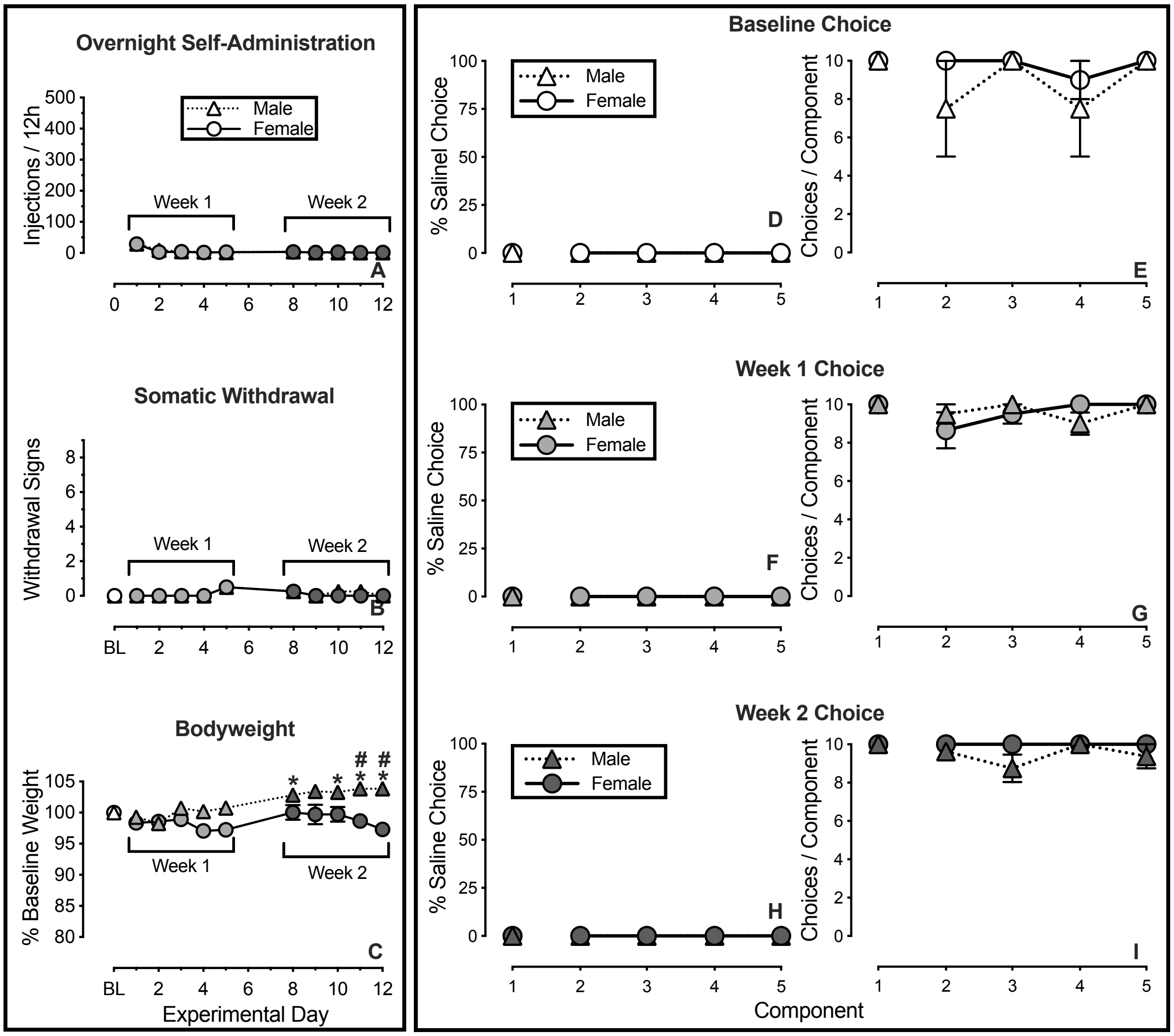

Supplemental Figure 2.

**Effects of overnight saline self-administration on saline-versus-food choice in male and female rats used in RNA sequencing studies.** Left panels (A-C): Overnight (6PM-6AM) saline self-administration and its effects on somatic withdrawal sign expression and bodyweight in male (n=4, left) and female (n=4, right) rats. Abscissae of left panels (A-C): experimental day. Top ordinate (A): number of saline infusions during each 12-h session. Middle ordinate (B): number of somatic withdrawal signs (maximum of nine) observed 8h after the conclusion of each overnight self-administration session. Bottom ordinate (C) change in bodyweight expressed as a percentage of baseline. Right panels (D-I): Sex comparison of saline-withdrawal effects on saline-versus-food choice assessed 8h after overnight (6PM-6AM) self-administration. Abscissae: component of the choice session. Left ordinates (D, F, H): percentage of completed ratio requirements on the saline-associated lever. Right ordinates (E, G, I): number of choices completed per component. Points represent mean ± SEM. * Denotes significant difference relative to Day 1 (A) or Baseline (B-C), which are placed above the symbol for female rats and below the symbol for female rats. # Denotes a significant sex difference at a particular experimental day. Significance defined as $p < 0.05$. See *Supplemental Table 1* for statistics relevant to each panel.

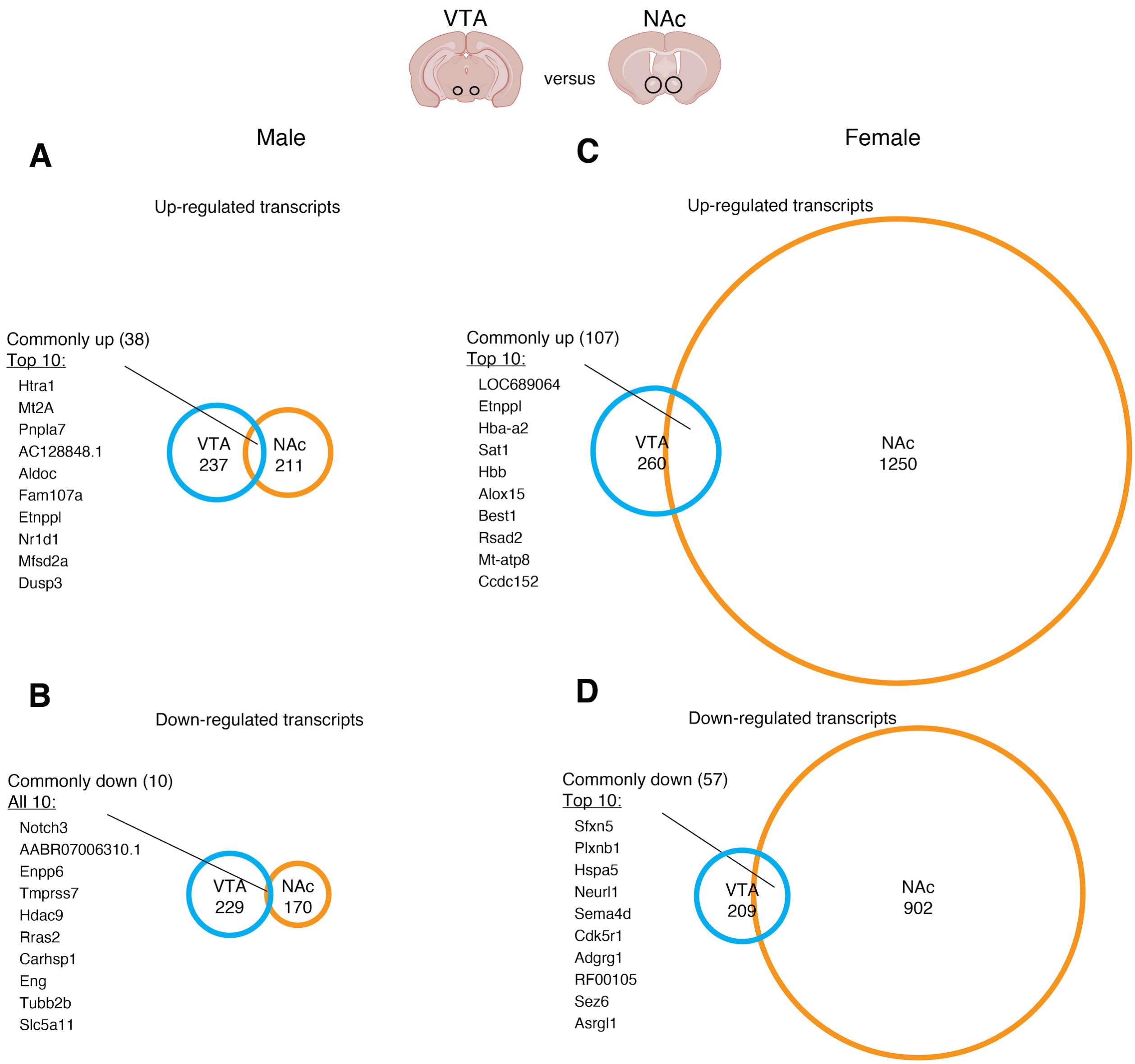

Supplemental Figure 3.

**Intra-sex comparisons of transcription across mesolimbic brain areas.**

Comparison of up- (A) and down-regulated (B) transcripts across the VTA and NAc of male mice. Comparison of up- (C) and down-regulated (D) transcripts across the VTA and NAc of female mice.

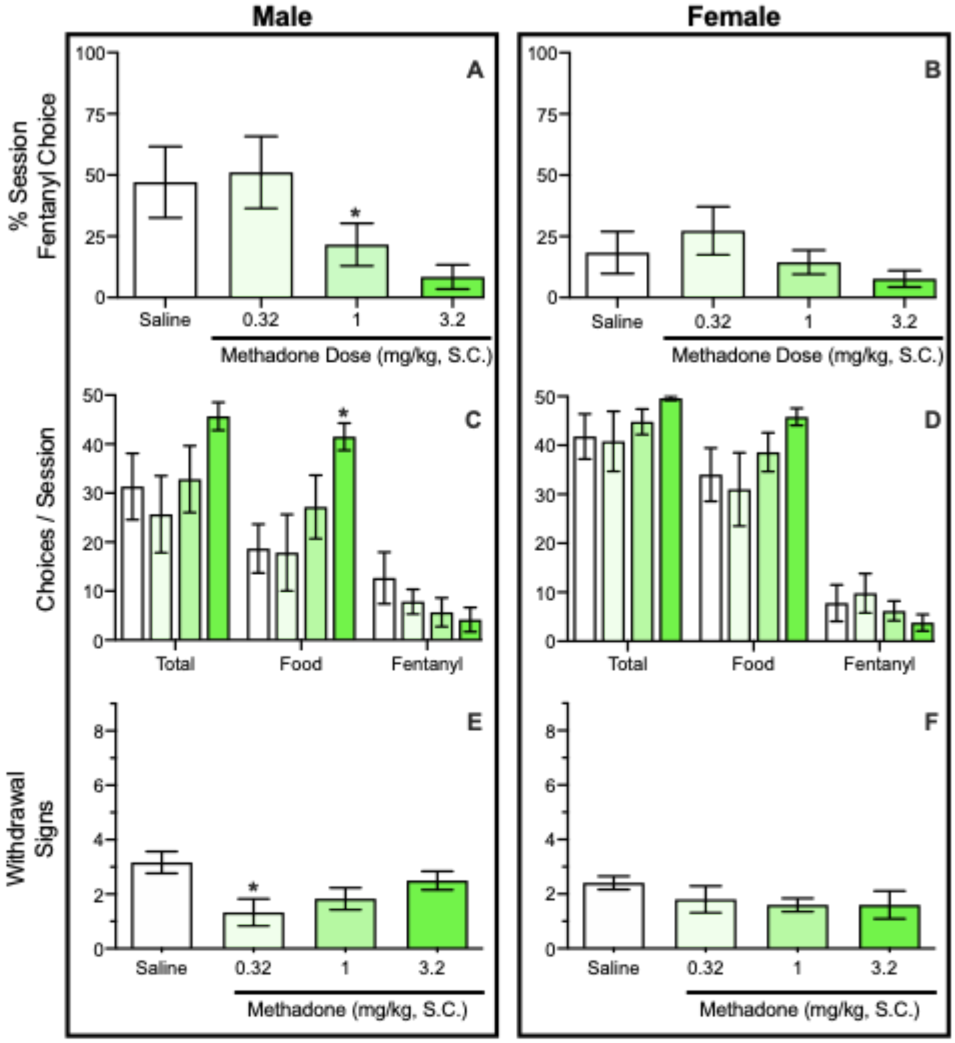

Supplemental Figure 4.

**Effects of acute methadone on fentanyl-versus-food choice and somatic withdrawal signs in opioid dependent male and female rats.** Effects of acute methadone treatment on fentanyl choice in male (n=6, left) and female (n=5, right) rats during Week 3 of overnight (6PM-6AM) fentanyl self-administration. Top and bottom abscissae: methadone dose. Middle abscissae: reinforcer type. Top ordinate (A-B): percentage of completed ratio requirements on the fentanyl-associated lever across the entire session. Middle ordinate: (C-D): number of choices completed per session. Bottom ordinate: (E-F): number of withdrawal signs present. Points represent mean ± SEM. *Denotes difference from saline. Significance defined as $p < 0.05$.
